# Emergence of the cortical encoding of phonetic features in the first year of life

**DOI:** 10.1101/2022.10.11.511716

**Authors:** Giovanni M. Di Liberto, Adam Attaheri, Giorgia Cantisani, Richard B. Reilly, Áine Ní Choisdealbha, Sinead Rocha, Perrine Brusini, Usha Goswami

## Abstract

Even prior to producing their first words, infants are developing a sophisticated speech processing system, with robust word recognition present by 4-6 months of age. These emergent linguistic skills, observed with behavioural investigations, are likely to rely on increasingly sophisticated neural underpinnings. The infant brain is known to robustly track the speech envelope, however to date no cortical tracking study could investigate the emergence of phonetic feature encoding. Here we utilise temporal response functions computed from electrophysiological responses to nursery rhymes to investigate the cortical encoding of phonetic features in a longitudinal cohort of infants when aged 4, 7 and 11 months, as well as adults. The analyses reveal an increasingly detailed and acoustically-invariant phonetic encoding over the first year of life, providing the first direct evidence that the pre-verbal human cortex learns phonetic categories. By 11 months of age, however, infants still did not exhibit adult-like encoding.

---

The human ability to understand speech relies on a complex neural system, whose foundations develop over the first few years of life. A wealth of evidence on the developmental progression of speech perception is available from infant behavioural studies, including with neonates, augmented by studies of speech production from around the second year of life^1,2^. Yet our understanding of speech perception in the first year of life is largely dependent on tasks relying on simple behaviours (e.g., head turn preference procedure). Direct investigation of the neural encoding of phonetic information in continuous natural speech across the first year of life has not previously been possible. Experiments using behavioural measures enable the assessment of valuable factors such as the familiarity of a particular speaker, the phonetic features that can be discriminated, and sensitivity to native versus non-native speech contrasts, thereby providing a time-line for the development of speech perception in the first year of life^1^. However, behavioural methods can only serve as an indirect index of the emergence of linguistic skills, and cannot reveal when the phonetic encoding in the human cortex becomes invariant across different instantiations. Previous behavioural studies focused on sound discrimination due to methodological constraints, and made use of targeted experimental paradigms involving simple stimuli. Although this behavioural timeline has been complemented by neurophysiological investigations, these studies have employed similar targeted paradigms, with the most widely-used neurophysiological measure with infants being the mismatch negativity (MMN, or mismatch response, MMR). The MMR is a neurophysiological signature of automatic change detection^3–5^ typically used to measure the ability to discriminate particular speech contrasts. However, previous studies showed that such mismatch responses in infants can sometimes be positive^6^, causing inconsistencies that can complicate or limit their use in infants. This leaves us with a number of key open questions: 1) How do infants perceive and encode the phonological units such as syllables and phonemes in **continuous natural speech?** 2) How are these speech sounds **encoded** in the infant brain? And 3) how does that encoding **develop across the first year of life?**

This study is the first to address these research questions directly. Non-invasive electroencephalography signals (EEG) were recorded as infants listened to 18 nursery rhymes (vocals only with no instruments involved) through video recordings of a native English speaker. EEG recordings were carried out at 4, 7 and 11 months of age from the first 50 participants in a longitudinal cohort involving 122 infants (the same subjects were tested in the three subsequent sessions and only participants with all sessions were selected). Three participants were excluded due to excessive EEG noise (see **Methods**). We then measured how the infant brain encodes acoustic and phonetic information by means of the multivariate Temporal Response Function analysis (TRF), a neurophysiology framework enabling the study of how neural signals encode continuous sensory stimuli^7,8^. TRF analyses were also carried out on recordings from adult participants listening to the same stimuli. We targeted one key aspect for speech perception, the perception of phonetic features. We do not assume here that encoding phonetic features equates to encoding phonemes, as there is a large psychoacoustic and developmental literature showing that phonemes are only represented by literate brains^9,10^. Our core hypothesis was rather that phonetic feature encoding (invariant to acoustic changes) would emerge in the neural responses to natural speech during the first year of life.

Speech TRFs reflect the neural tracking of (or neural entrainment to, in the broad sense^11^) natural speech features (e.g., acoustic envelope), offering a direct window into human perception during natural listening without imposing any particular task other than listening. In recent years, neural tracking measures have played a growing role in the study of speech comprehension and auditory processing in general. Many TRF studies have assessed the neural tracking of the acoustic envelope^12–15^, which is an important property of speech that co-varies with a number of key properties of interest (e.g., syllable stress patterns, syllables, phonemes). Neural tracking of the speech envelope (or envelope tracking) was shown to reflect both bottom-up and top-down cortical processes in adult listeners, encompassing fundamental functions such as selective attention^15–17^, working memory processing load^18^, and prediction^19,20^. While robust envelope tracking has also been demonstrated in infants^21–26^, envelope measures only reveal some of the cortical mechanisms underlying speech perception. Recent work with adults and children has demonstrated that TRFs can be extended to isolate the neural encoding of targeted speech properties of interest, starting from phonetic features^27^. Phonetic encoding was measured in multiple studies from different research teams^27–31^, and the neural tracking was shown to correlate with phonemic awareness skills in school-aged children between 6 and 12 years of age^32^ and with second language proficiency in adults^33^.

Here, we employed TRFs to test the hypothesis that the neural encoding of phonetic features during natural speech listening is already developing during the first-year of life. Current behavioural data indicate that infant perception becomes more selective towards native than non-native speech contrasts around 9-12 months of age^34^ (see footnote^i^), with perceptual “magnet” effects helping to isolate native from non-native phonetic contrasts already by 6 months^35^. We hypothesised that these phenomena may be underpinned by a progressively more precise and acoustically-invariant neural encoding of phonetic features across the first year of life. This encoding would be expected to emerge as a neural response to speech that reflects a growing invariance towards phonetic categories, where the limit case would be to have neural responses to phonetic categories that are fully invariant to acoustic changes. This longitudinal investigation offers the first view into phonetic feature encoding in the first year of life, while accounting for the full complexity of speech in naturalistic listening environments. In the discussion, we connect our encoding analyses with previous work on phonetic discrimination and consider the key role of perceptual invariance. Our results provide a promising new avenue for developmental research with both infants and children. Precise measures of how and when phonetic feature encoding evolves could serve as a complementary set of risk factors for developmental language disorders, as well as illuminating the phonological trajectories experienced by both typically- and atypically-developing children.

## Results

### Robust neural tracking of acoustic and phonetic features in infants

A multivariate TRF analysis was carried out to assess the low-frequency (1-15 Hz) neural encoding of speech across the first year of life. Acoustic and phonetic features were extracted from the stimulus. Acoustic features consisted of the 8-band acoustic spectrogram of speech (*S*) sound and the half-way rectified envelope derivative (*D*). Fourteen phonetic features were included to mark the categorical occurrence of speech sounds, according to articulatory features describing voicing as well as manner and place of articulation. To account for possible differences in the encoding of stressed and unstressed sounds, each phonetic feature produced to two distinct vectors, leading to a 28-dimensional phonetic features matrix (*F*; see **Methods**). A nuisance regressor was also included to capture EEG variance related to visual motion (*V*). Single-subject TRFs were derived for each experimental session to assess the cortical encoding of acoustic and phonetic features by fitting a multivariate lagged regression model with all such features simultaneously (**Figure 1A**).

**Figure 1:**
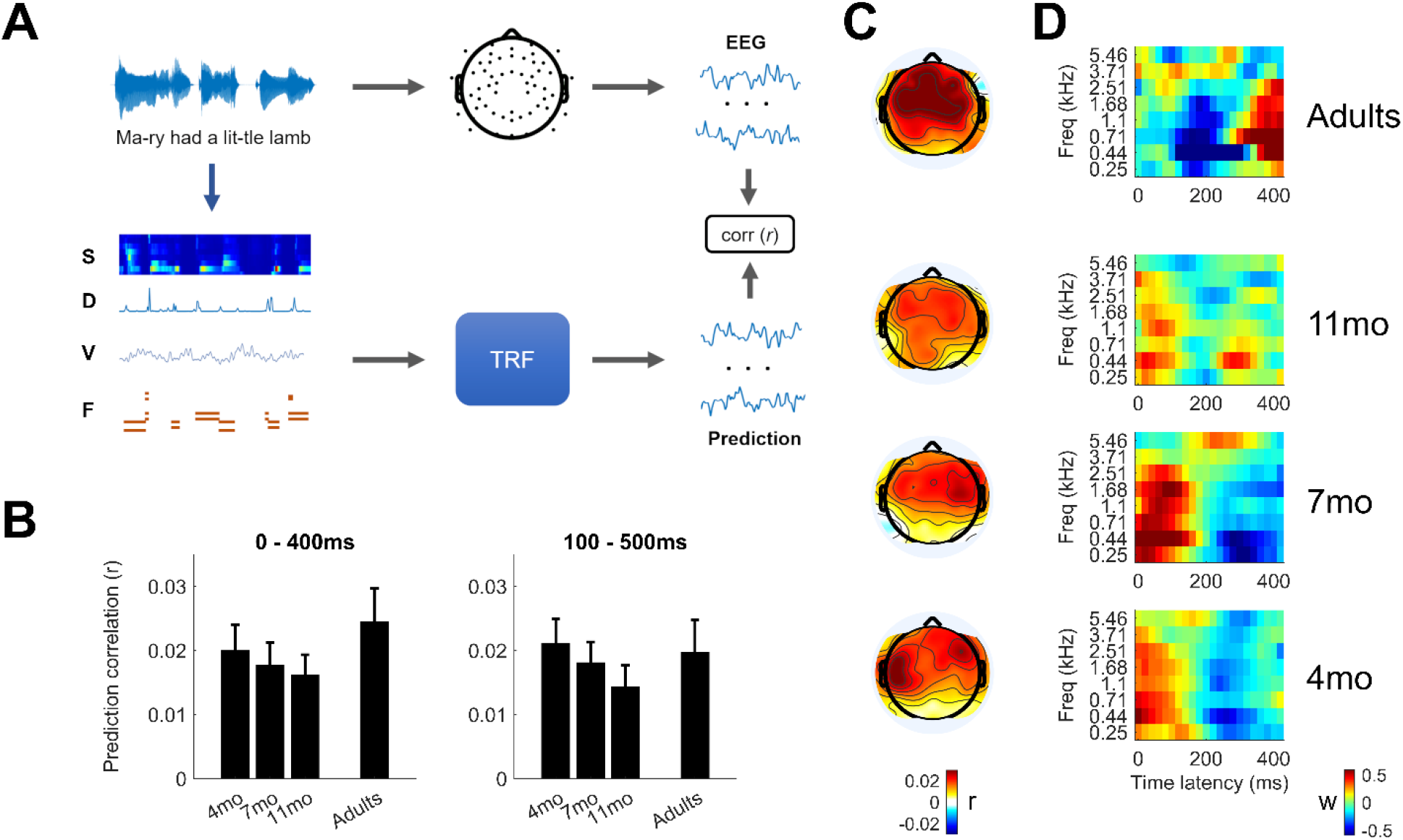
EEG tracking of acoustic and phonetic features in infants and adults. **(A)** Schematic diagram of the analysis paradigm. Multivariate Temporal Response Function (TRF) models were fit to describe the forward relationship between speech features and the EEG signal recorded from adults and infants (4, 7, and 11mo). Speech features included the 8-band acoustic spectrogram (S), half-way rectified envelope derivative (D), visual motion (V), and phonetic features (F). **(B)** EEG prediction correlations of the multivariate TRF model were significant within each group for both the time-lag windows 0-400ms and 100-500ms. **(C)** Topographical patterns of the EEG prediction correlations in infants (shown for the TRF window 0-400ms) became progressively more similar to adults responses with age. **(D)** TRF weights corresponding to the S features averaged across centro-frontal electrodes.

EEG prediction correlations, calculated with leave-one-out cross-validation and averaged across all EEG channels, were **greater than zero for all age groups** (one-sample Wilcoxon rank sum test, FDR-corrected for multiple comparisons; 4mo: *p*=9.5*10^-6^; 7mo: *p*=9.5*10^-6^; 11mo: *p*=9.5*10^-6^; adults: *p*=2.2*10^-4^; **Figure 1B**). Consistent with previous work, this analysis was carried out by considering speech-EEG lags from 0 to 400ms, which were shown to largely capture the cortical acoustic-phonetic response in adults. The TRF analysis was also repeated when considering a 100-500ms lag window, aiming to control for possible responses with longer latencies in infants, while keeping the same model complexity (i.e., same window size). This analysis also led to significant EEG predictions (one-sample Wilcoxon rank sum test, FDR-corrected; 4mo: *p*=3.6*10^-6^; 7mo: *p*=3.6*10^-6^; 11mo: *p*=1.7*10^-6^; adults: *p*=0.001; **Figure 1B**), indicating a consistent speech-EEG relationship involving acoustic and phonetic features in both time windows.

Topographic differences were expected both across participants and by age group due to major anatomical changes during infancy^36^. Larger EEG prediction correlations were measured in centro-frontal electrodes for all age groups (**Figure 1C**), with **topographies becoming progressively more similar to those for adults** with age in both the 0-400ms lag window (bootstrap with group size = 17 and 1000 iterations; average correlation with adults: *r* = 0.43, 0.51, 0.54 for 4mo, 7mo, and 11mo respectively; repeated measures ANOVA on infant data with age as the repeated factor: *F*(2,1998) = 172.8, p = 6.1*10^-70^) and 100-500ms window (bootstrap; average correlation with adults: *r* = 0.32, 0.55, 0.52 for 4mo, 7mo, and 11mo respectively; repeated measures ANOVA: *F*(2,1998) = 900.6, p = 1.5*10^-279^). TRF models corresponding to spectrogram features are reported in **Figure 1D**, where weights were averaged across fourteen centro-frontal electrodes (25% of all channels) and all participants (see **Methods**).

### Emergence of phonetic feature encoding in the first year of life

The analyses that follow aim to determine if and when cortical signals encode acoustically-invariant phonetic features during the first year of life. In line with previous behavioural work^35,37–41^ and current developmental theories^1,34^, we expected categorical phonetic feature encoding to emerge from 6 months on (i.e., from the 7mo recording session, in the present study), with progressively stronger encoding across the first year of life visible by 11 months of age. To test this hypothesis (see Hp2 in **Figure 2A**), phonetic feature encoding was assessed based on the multivariate TRF models described in the previous section. Neural activity linearly reflecting phonetic feature categories but not sound acoustics was accounted by subtracting EEG prediction correlations corresponding to acoustic-only TRFs (which did not include phonetic features; see **Methods**) from those corresponding to acoustic-phonetic TRFs^ii^. For consistency with previous work^27,29,32,33,42–44^, this metric is referred to as FS-S (F: phonetic features; S: spectrogram and envelope derivative). As expected (Hp1-3), FS-S values were progressively larger with age (average across all EEG channels; **Figure 2B)**, with values greater than zero emerging from 11 months of age for the time-latency window 0-400ms (one-sample Wilcoxon rank sum test, FDR-corrected; 4mo: *p*=0.237; 7mo: *p*=0.237; **11mo: *p*=0.037**; **adults: *p*=0.037**; black bars indicate *p*<0.05), and **from 7 months** of age for the time-latency window 100-500ms (one-sample Wilcoxon rank sum test, FDR-corrected; 4mo: *p*=0.167; **7mo: *p*=0.044**; **11mo: *p*=0.023**; **adults: *p*=0.044**). This latter finding is consistent with hypothesis 2 (Hp_2_), as depicted in **Figure 2A**. Furthermore, the 4mo group did not show significant FS-S values for subsequent latency windows (200-600ms and 300-700ms).

**Figure 2:**
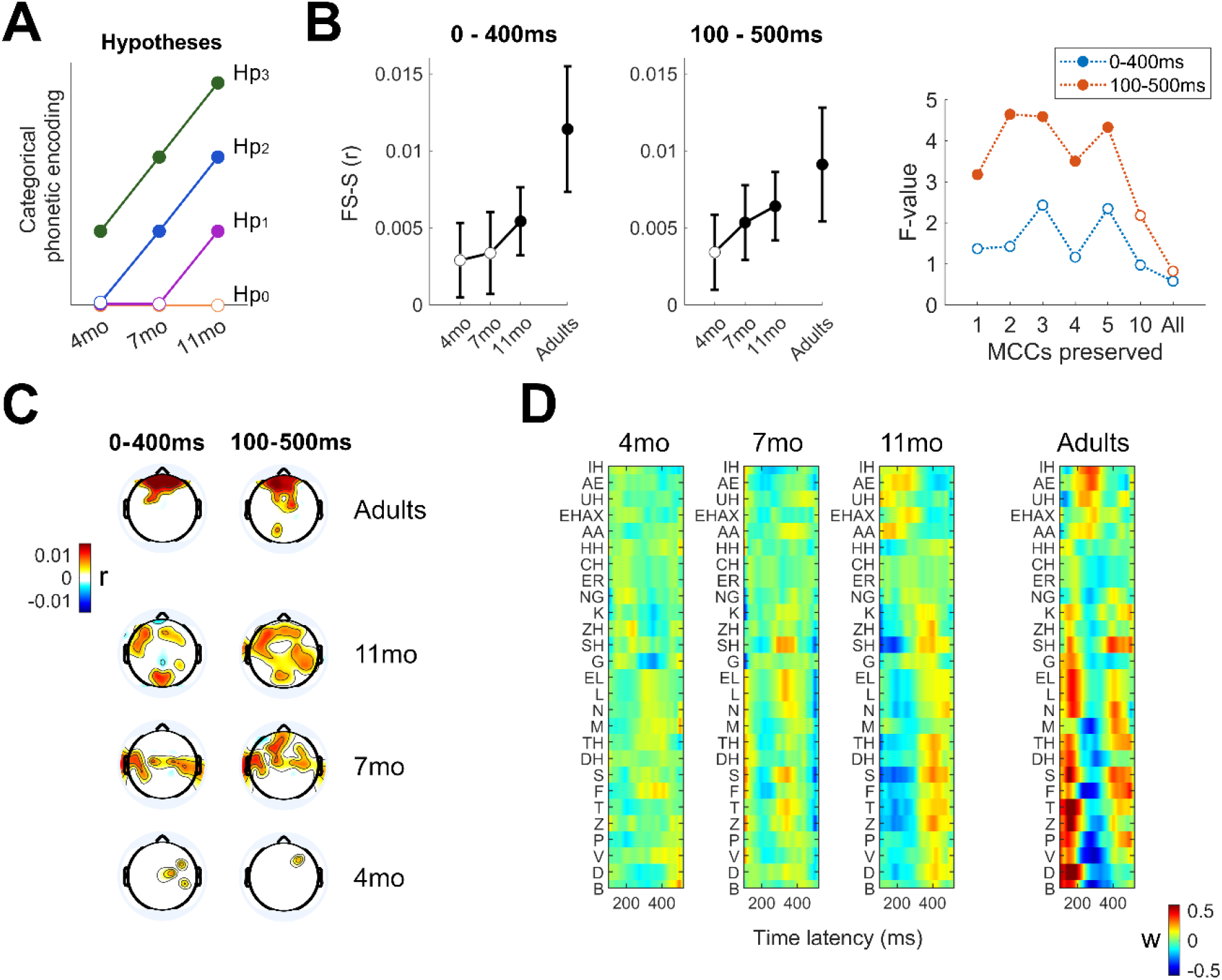
Cortical encoding of phonetic features in the first year of life. **(A)** Hypotheses: The cortical encoding of phonetic feature categories was expected to emerge and progressively increase across the first year of life. Hypothesis 0 (Hp0): No phonetic encoding in the first year of life; Hp1-3: phonetic encoding from 11, 7, and 4 months of age respectively. **(B)** Phonetic feature encoding measured as the EEG prediction correlation gain when including phonetic features in the TRF (mean and SE across participants, for the 0-400ms and 100-500ms lag windows). Black bars indicate significance (p<0.05 after FDR-correction). The right panel indicates the F-statistics (repeated measures ANOVA) when using MCCA denoising (retaining 1, 2, 3, 4, 5, and 10 components) and without MCCA denoising (‘all’). A main effect of age emerged for the 100-500ms TRF when retaining up to 5 components (filled dots indicate significance; p<0.05). **(C)** Phonetic feature encoding (EEG prediction correlation gain) across all electrodes. Coloured areas indicate significance (p<0.05, t-test with FDR correction). **(D)** TRF weights corresponding to phonetic features for the 100-500ms TRF.

**Figure 2:**
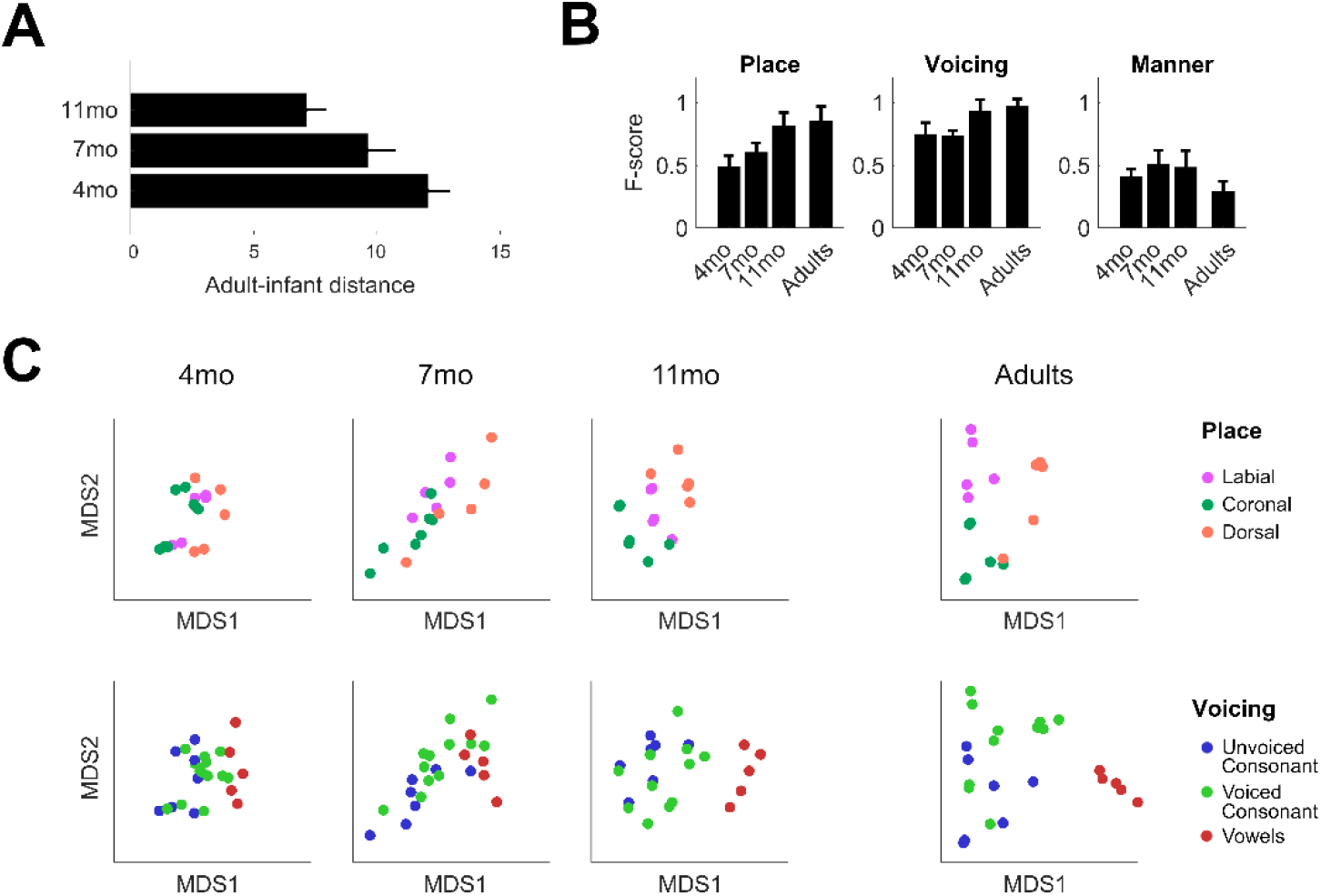
Sensitivity to phonetic feature groups in the first year of life. **(A)** Distance between infant and adult TRF weights (mean and SE). **(B,C)** Multidimensional scaling maps (MDS) were calculated on the phonetic features TRFs as a function of peri-stimulus time lag and electrode. By carrying out 100 repeated k-means classification, F-score measures were derived representing the discriminability of specific phonetic feature groups in the TRFs (mean and SE across repetitions) i.e., place of articulation, voicing, and manner of articulation (B). Individual MDS maps are shown in (C), where dots correspond to adult phonemes.

While the previous analysis identified significant phonetic encoding within individual age groups, the analysis that follows explicitly assessed if phonetic encoding increased across the first year of life. TRF models and the corresponding EEG prediction correlations showed large between-subject variability, which was expected due to the noisy single-subject data. Under the assumption that participants in the same age group present EEG responses to nursery rhymes with similar temporal patterns, a Multiway Canonical Correlation Analysis (MCCA)^45^ was carried out to isolate EEG components that are consistent within each group, substantially improving the signal-to-noise ratio of the single-subject EEG. A repeated measures ANOVA test was then carried out to determine if FS-S increased with age by considering the first MCCA component only (MCC1) i.e., the EEG component with highest temporal correlation across subjects within a given age group. A significant increasing trend emerged for the 100-500ms window (*F*(2,138)=3.19, *p*=0.044) but not for the 0-400ms latency window (*F*(2,138)=1.37, *p*=0.257; see coloured panel **Figure 2B**). The test was also run when considering an increasing number of MCCs, showing significant results when considering up to five components for the 100-500ms latency window. The EEG encoding of phonetic features was also studied at individual electrodes, revealing robust encoding of phonetic features on large clusters of EEG channels in adults as well as infants from 7 months of age, both when considering 0-400ms and 100-500ms windows (**Figure 2C**; FDR-corrected one-sample Wilcoxon rank sum tests were run on each EEG channel; colours indicate significant results with *p*<0.05).

Further analyses were carried out to assess the phonetic feature encoding at a fine-grained level, by studying the TRF weights of the acoustic-phonetic TRF (weights are shown in **Figure 2D**). Phonetic distance maps^33^ were calculated by using a multidimensional scaling analysis (MDS; see **Methods**) on the TRF weights corresponding to phonetic features. This approach allows to quantify and visualise the level of similarity or distance between datapoints by accounting simultaneously for multiple EEG channels and peri-stimulus time latencies. By selecting the two most relevant MDS dimensions, the infant-adult Euclidean distance was calculated for each age group (**Figure 3A**; bars indicate mean and SE calculated across 27 adult phonemes computed as a linear combination of the corresponding phonetic features), showing that **the infant-adult distance decreases with infant age** in the first year of life (repeated measures ANOVA, *F*(2,52) = 18.2, *p* = 5.0*10^-7^). *F*-score measures were derived quantifying the discriminability of specific phonetic feature groups in the individual-subject TRFs using a *k*-means analysis (mean and SE were calculated on the *F*-scores resulting from the 100 repetitions of *k*-means). The phonetic feature groupings considered for this analysis were place of articulation, voicing, and manner of articulation (**Figure 3B,C**). As a validation step, the stability of the resulting *F*-scores for the infants in the three longitudinal sessions was assessed over 100 repetitions of the *£*-means procedure (repeated measures ANOVA, voicing: *F*(2,198) = 155.8, *p* < 10^-12^; place of articulation: *F*(2,198) = 377.2, *p* < 10^-12^; manner of articulation: *F*(2,198) = 29.4, *p* = 6.84*10^-12^; **Figure 3B**). Next, statistical analyses were carried out to determine the significance of the result across participants. As expected (due to factors such as low-SNR, limited data, and inter-subject variability), single-subject phonetic feature maps did not lead to significant results, even though results for place of articulation were trending towards significance (repeated measures ANOVA, voicing: *F*(2,92) = 0.6, *p* = 0.54; place of articulation: *F*(2,92) = 2.6, *p* = 0.08; manner of articulation: *F*(2,92) = 0.5, *p* = 0.61). To compensate for the limited single-subject data, we ran a bootstrap analysis with 100 repetitions, each derived by averaging 17 subjects i.e., the same number of participants in the adult group. This analysis revealed that phonetic feature encoding increased with age for place of articulation and voicing, but not manner of articulation (repeated measures ANOVA, **voicing**: *F*(2,198) = 21.3, ***p* = 4.2*10^-9^**; **place of articulation**: *F*(2,198) = 17.6, ***p* = 9.0*10^-8^**; manner of articulation: *F*(2,198) = 1.5, *p* = 0.22).

## Discussion

The present investigation offers the first direct evidence that the human cortex encodes phonetic categories during the first year of life, demonstrating significant phonetic encoding from 7 months of age and progressively stronger encoding thereafter. A fine-grained and longitudinal understanding of the development of phonetic feature encoding by the same infants listening to continuous speech was previously absent from the literature. The behavioural and MMR infant speech processing literature has used targeted experimental contrasts, focused largely on the perception of syllable stress and speech rhythm and on phonetic category formation. As rhythm and stress patterns aid in identifying word boundaries, and phonetic categories aid in comprehension (e.g., distinguishing ‘doggy’ from ‘daddy’), this prior work has been important, showing that infants are sensitive to differences in speech rhythm from birth^46,47^, and are sensitive to some phonetic information as neonates^48^. Nevertheless, no prior study has used continuous speech as a basis for studying phonetic encoding. Consequently, our findings have several implications for understanding of the development of speech processing.

Currently, it remains unclear how and at what stage of development phonetic category encoding is learnt. This question remains open largely because of methodological constraints. The present study offers, for the first time, direct evidence on the ‘when’ of phonetic category learning. There is a consensus in the literature that discriminating phonetic categories is a key processing step regarding speech comprehension by adults^49^, although see Feldman et al., 2021^50^ for recent caveats regarding infants. While adult studies used direct invasive recordings to measure the cortical encoding of phonetic categories^51^, recent methodological developments (i.e., the TRF framework^7,8^) allowed us to circumvent some of the major challenges encountered by previous infant studies, thereby providing more precise developmental information.

The assessment of phonetic encoding as operationalised here fulfils three main elements of novelty that go beyond any previous investigation. First, we studied the cortical encoding of phonetic categories in infants with **direct neural measurements** based on EEG and as part of an unprecedented targeted longitudinal investigation. Second, the use of the forward TRF framework allowed us to assess phonetic category **encoding**, rather than relying on the typical sound discrimination metrics used in prior behavioural^35,37–41^ and neurophysiology studies (e.g., MMR)^4,52–56^. Third, the TRF framework allowed us to study the perception of **natural speech** in infants, instead of focusing on selected phonetic or word contrasts, as in the past literature. This is a crucial step forward, as the discriminatory skills that infants exhibit in simplified laboratory settings (e.g., isolated syllable discrimination measured via a head-turn or looking procedure or with MMRs) may not be sufficient for detecting phonetic categories in naturalistic settings.

The present study indicates that phonetic category encoding during natural speech listening emerges between 5 and 7 months of age. This provides the literature with new and fundamental insights into the development of speech processing in neurotypical infants. Further, the TRF approach yields novel information on *which* specific phonetic contrasts evolve with age, demonstrating a natural progression toward adult phonetic encoding. This insight is further reinforced by the observation that acoustic encoding did not increase with age. Consequently, the enhanced phonetic encoding with age observed here could not simply be due to stronger acoustic encoding, as acoustic encoding showed the opposite pattern (a non-significant decreasing trend with age). Interestingly, our results suggest that 4mo pre-babbling infants, despite being equipped with the fundamental combinatorial code for speech analysis^57^, do not yet exhibit categorical phonetic encoding. Based on these results, we can speculate that prior demonstrations of infant behavioural and MMR discrimination between syllables like “pa” and “ba” probably have an acoustic basis but do not reflect categorical phonetic encoding. In other words, the ability to distinguish two sounds does not necessarily mean that those sounds are encoded as separate categories. Our study is instead probing that categorical encoding directly.

One challenge with longitudinal neurophysiology studies in infants is the substantial anatomical change that occurs with age, meaning that while macroscopic patterns are likely to remain consistent (e.g., temporal vs. occipital), there cannot be a channel-by-channel correspondence between age groups, even when considering the same participants. For this reason, the majority of this investigation focused on measures combining multiple EEG channels simultaneously (e.g., **Figure 1D** was an average of 14 centro-frontal channels). These considerations make the topographical distribution of phonetic encoding strength shown in **Figure 2C** even more remarkable, as a centro-frontal cluster of EEG channels was shown to reflect phonetic encoding across all age groups with the exception of 4 months, where phonetic encoding as assessed by the TRF was not significant.

Our phonetic encoding results showed topographical patterns and TRF weights for adults that differ from the prior adult EEG literature on natural speech listening TRFs^27,29^. While part of the discrepancy may be due to the use of a different EEG acquisition device and to the use of audio-visual stimuli, the primary explanation is likely to be the choice of stimuli. This is the first TRF investigation of phonetic processing with adults involving a nursery rhyme listening task. Nursery rhymes are indeed a form of natural speech which is more suited to infants. The rhythmic cues and exaggerated stress patterns characterising nursery rhymes have been demonstrated to be important elements supporting speech perception and language learning^58,59^, accordingly they were ideal stimuli for the Cambridge UK BabyRhythm study. In prior TRF work, we have demonstrated similar envelope entrainment to these nursery rhymes by adults and infants^26^. Nevertheless, it is important to note that the regular rhythms and melodic properties of nursery rhymes makes the different from the typical speech TRF stimuli used with adults, such as audio-books and podcasts. As such, the TRF results were expected to show different spatio-temporal patterns for adult listeners compared to previous TRF work.

The results of this study add to the growing literature on cortical speech tracking^21,27,31,33,44,60–62^. While the literature typically focuses on the cortical tracking of the speech envelope^16,24,63–66^ (including previous analyses of this dataset^21,26^), the present investigation enriches our understanding of phonetic feature TRFs. Prior TRF studies of phonetic encoding in adults and children have revealed that phonetic processing is affected by speech clarity^43^, selective attention^29^, and proficiency in a second language^33^, and shows correlations with psychometric measures of phonemic awareness^32^. The present study demonstrates that emergent phonetic TRFs can also be measured in pre-verbal infants, providing a novel window into infant perception and cognition. Whilst recent developments have started to use neural tracking to predict language development in infants^67^, further research will also determine whether a robust relationship exists between speech TRFs and other related aspects of cognition (e.g., selective attention, prediction) in infants, and when such related aspects come on-line. Further research with infants at family risk for disorders of language learning may also reveal when and how developmental trajectories are impacted by developmental disorders that are carried genetically, such as developmental dyslexia and developmental language disorder. Such work could be very valuable regarding early detection and improved mechanistic understanding of these disorders.

In summary, this study demonstrated the emergence of phonetic encoding from 7 months of age using direct neural measurements during natural speech listening. The data provide clear-cut evidence of the emergence of phonetic categories that contributes to the current debate regarding their role in the development of speech processing. Our demonstration that phonetic encoding can be assessed with nursery rhyme stimuli in ecologically-valid conditions opens the door to cross-language work using TRFs that investigates the interaction between characteristics of natural language such as phonological complexity and the development of phonetic encoding. It also provides opportunities for novel mechanistic investigations of the development of bi-lingual and multi-lingual lexicons during language acquisition.

## Online Methods

### Subjects and experimental procedure

The present study carried out a re-analysis of an EEG dataset involving a speech listening task in a longitudinal cohort of fifty infants (first part of a larger cohort of 122 subjects^21^). Participants were infants born full term (37-42 gestational weeks) and had no diagnosed developmental disorder, recruited from a medium sized city in the United Kingdom and surrounding areas via multiple means (e.g., flyers in hospitals, schools, and antenatal classes, research presentations at maternity classes, online advertising). The study was approved by the Psychology Research Ethics Committee of the University of Cambridge. Parents gave written informed consent after a detailed explanation of the study and families were repeatedly reminded that they could withdraw from the study at any point during the repeated appointment. The experiment involved three EEG recording sessions when the infants (24 male and 26 female) were 4 months old (4mo; 115.6 ± 5.3 days), 7 months old (7mo; 212.5 ± 7.2 days) and 11 months old (11mo; 333.0 ± 5.5 days) [mean ± standard deviation (*SD*)]. A bilingualism questionnaire (collected from 45 out of the 50 infants) ascertained that 38 of the infants were exposed to a monolingual environment and 12 were exposed multilingual environment, of these 93.5% (43 infants) reported English as the primary language exposed to the infant. Note that this was a longitudinal investigation, meaning that the same 50 infants were tested at 4, 7, and 11 months of age. In addition to the 150 EEG sessions from the infant dataset, this study also analysed EEG data from twenty-two monolingual, English-speaking adult participants performing the same listening task (11 male, aged 18-30, mean age: 21). Data from four adult participant was excluded due to inconsistencies with the synchronisation triggers, leaving seventeen participants data for the analysis.

Infant participants were seated in a highchair (one metre in front of their primary caregiver) in a sound-proof acoustic chamber, while adult participants were seated in a normal chair. All participants were seated 650mm away from the presentation screen. EEG data were recorded at a sampling rate of 1 kHz using a GES 300 amplifier using a Geodesic Sensor Net (Electrical Geodesics Inc., Eugene, OR, United States). 64 and 128 channels were used for infants and adults respectively. Sounds were presented at 60 dB from speakers placed either side of the screen (Q acoustics 2020i driven by a Cambridge Audio Topaz AM5 Stereo amplifier). Participants were presented with eighteen nursery rhyme videos played sequentially, each repeated 3 times (54 videos with a presentation time of 20’ 33’’ in total). Adult participants were asked to attend to the audio-visual stimulus while minimising their motor movements. All adult participants completed the full experiment. Infants listened to at least two repetitions of each nursery rhyme (minimum of 36 nursery rhymes lasting 13’ 42’’). The experiment included other elements that were not relevant to the present study (e.g., resting state EEG; please refer to the previous papers on this dataset for further information^21,26^).

### Stimuli

A selection of eighteen typical English language nursery rhymes was chosen as the stimuli. Audio-visual stimuli of a singing person (upper-body only) were recorded using a Canon XA20 video camera at 1080p, 50fps and with audio at 4800 Hz. A native female speaker of British English used infant-directed speech to melodically sing (for example “Mary Quite Contrary”) or rhythmically chant (for nursery rhymes like “There was an old woman who lived in a shoe”) the nursery rhymes whilst listening to a 120 bpm metronome through an intra-auricular headphone (e.g., allowing for 1Hz and 2Hz beat rates; see Figs. S2 and S4 from Attaheri et al.^21^). The metronome’s beat was not present on the stimulus audios and videos, but it ensured that a consistent rhythmic production was maintained throughout the 18 nursery rhymes. To ensure natural vocalisations, the nursery rhyme videos were recorded sung, or rhythmically chanted, live to an alert infant.

### Data preprocessing

Analyses were conducted with MATLAB 2021a by using custom scripts developed starting from publicly available scripts shared by the CNSP initiative (Cognition and Natural Sensory Processing; https://cnspworkshop.net; see section Data and Code Availability for further details).

In order to carry out the same preprocessing and analysis pipeline on infants and adult EEG data, the adult 128-channel EEG data was transformed into a 64-channel dataset via spline interpolation, with the relative channel locations corresponding to those of the infant participants. All subsequent analyses on infants and adult were identical.

The four facial electrodes (channels 61-64) were excluded from all analyses, as they are not part of the specific infant-sized EGI Geodesic sensor net. The EEG data from the remaining 60 channels was band-pass filtered between 1 and 15 Hz by means of zero-phase shift Butterworth filters with order 2 (by using the filtering functions in the CNSP resources). EEG signals were downsampled to 50 Hz. Next, Artifact Subspace Reconstruction (ASR; *clean_asr* function from EEGLAB^68^) was used to clean noise artifacts from the EEG signals. Channels with excessive noise (which could not be corrected with ASR) were identified via probability and kurtosis and were interpolated via spherical interpolation, if they were three standard deviations away from the mean. EEG signals were then re-referenced to the average of the two mastoid channels, which were then removed from the data, producing a preprocessed EEG dataset with 58 channels. Data from repeated trials was then averaged. Three infant subjects were removed because of excessive noise in at least one of their three recording sessions.

### Sung speech representations

The present study involved the measurement of the coupling between EEG data and various properties of the sung speech stimuli. These properties were extracted from the stimulus data based on methodologies developed in previous research. First, we defined a set of descriptors summarising low-level acoustic properties of the speech stimuli. Acoustic features consisted of an 8-band acoustic spectrogram (S) and a half-way rectified broadband envelope derivative (D)^33,60^. S was obtained by filtering the sound waveform into eight frequency bands between 250 and 8 kHz that were logarithmically spaced according to the Greenwood equation^69^. The broadband envelope was calculated as the sum across the eight frequency bands of S. The D signal was then derived by calculating the derivative of the broadband envelope, and by half-way rectifying the resulting signal. Second, fourteen phonetic features were then selected to mark the categorical occurrence of speech sounds, according to articulatory features describing voicing, manner, and place of articulation^70,71^: voiced consonant, unvoiced consonant, plosive, fricative, nasal, strident, labial, coronal, dorsal, anterior, front, back, high, low. To account for possible differences in the encoding of stressed and unstressed sounds, each phonetic feature was assigned to two distinct vectors, leading to a 28-dimensional phonetic features matrix (*F*). The precise timing of the phonetic units was identified in three steps. First, syllable and phoneme sequences were obtained from the transcripts of the nursery rhymes. Second, an initial alignment was derived by identifying the syllabic rate and syllable onsets for each piece, and then assigning the phonemes in a syllable starting from the corresponding onset time. This automatic alignment was stored according to the TextGrid format^72^. Third, the phoneme alignments were manually adjusted using Praat software^72^. Phonetic feature vectors were produced in MATLAB software to categorically mark the occurrence of phonetic units from start to finish with unit rectangular pulses^27^. Finally, a nuisance regressor was also included to capture EEG variance related with visual motion (V), which was derived as the frame-to-frame luminance change, averaged across all pixels.

### Multivariate Temporal Response Function (mTRF)

A single input event at time *t_0_* affects the neural signals for a certain time window [*t_1_, t_1_*+*t*_win_], with *t_1_* ≥ 0 and *t_win_* > 0. Temporal response functions (TRFs) describe this relationship at the level of individual subject and EEG channel. In this study, TRFs were estimated by means of a multivariate lagged regression, which determines the optimal linear transformation from stimulus features to EEG (forward model)^13,73^. A multivariate TRF model (mTRF) was fit for each subject by considering all features simultaneously (S, D, F, and V; **Figure 1A**) with the mTRF-Toolbox^7,8^. While a time-lag window of 0-400 ms was considered sufficient to largely capture the acoustic-phonetic/EEG relationship with a single-speaker listening task in adults, based on previous studies (e.g., Di Liberto et al.^27^), the relevant latencies in infants were unknown. To account for possible slower or delayed response in infants, a time-latency window of 100-500 ms (i.e., same duration but with longer latency) was also included in the analysis for all groups. The reliability of the TRF models was assessed using a leave-one-out cross-validation procedure (across trials i.e., nursery rhymes), which quantified the EEG prediction correlation (Pearson’s *r*) on unseen data while controlling for overfitting. The TRF model calculation included a Tikhonov regularization, which involves the tuning of a regularization parameter (λ) that was conducted by means of an exhaustive search of a logarithmic parameter space from 0.01 to 10^6^ on the training fold of each cross-validation iteration^7,8^. Note that the correlation values were calculated with the noisy EEG signal; therefore, the *r*-scores could be highly significant even though they have low absolute values (*r* ~ 0.1 for sensor-space low-frequency EEG^27,30,60^).

### Phonetic distance maps

We sought to study the effect of development on phonetic perception by projecting the TRF weights corresponding with the phonetic features onto a space in which distance represents the perceptual separation between phonological units. The TRF weights for the phonetic features, which were represented in a 28-dimensional space, were projected to the phoneme space. To do so, the TRF for a given phoneme was calculated as the sum of the TRF weights of the corresponding features. This produced a 54-dimensional matrix: 27 stressed and 27 unstressed phonemes. The weights corresponding to the two versions of a phoneme (stressed and unstressed) were then combined when projecting to the phonetic feature map, obtaining a 27-dimensional space. Specifically, a classical multidimensional scaling (MDS) was used to project the phoneme TRF weights (phonemes were considered as objects and time latencies were considered as dimensions) onto a multidimensional space for each age group, in which distances represented the discriminability of particular phonetic contrasts in the EEG signal. The result for each infant group was then mapped to the average adult MDS space by means of a Procrustes analysis (MATLAB function *procrustes*). This analysis allowed us to project the infant phonetic feature maps for different proficiency levels to a common multidimensional space where they could be compared quantitatively; we call these maps phonetic feature distance maps. Note that while this transformation does not assume nor imply categorical phonemic encoding, it indeed allows to quantify and visualise the encoding of phonetic features across age, by using familiar phonemic units.

While **Figure 3A** shows the infant-adult distances in the MDS space, the effect of age on phonetic feature encoding was also quantified with a clustering analysis. Specifically, the randomised clustering algorithm *k-* means was run on the MDS maps for each group to determine whether the phonetic feature groups could be deduced from the data without supervision. We performed 100 repetitions of *k*-means and a classification *F*-Score (or F_1_-Score) was calculated on each of them, obtained as the harmonic mean of precision and recall. We performed this procedure for the three feature sets of interest: a) the three-class feature-set with ‘vowels’, ‘voiced consonants’, and ‘unvoiced consonants’; b) the three-class feature-set describing the place of articulation, with features ‘labial’, ‘coronal’, and ‘dorsal’; and c) the three-class feature-set describing the manner of articulation, with features ‘fricative’, ‘stop’, and ‘nasal’. Since all feature-sets were three-dimensional, *k* had always value three. Note that *k*-means is an unsupervised clustering algorithm, meaning that there was no direct correspondence between the clusters and the classes of interest. As such, *F*-scores were selected for the best matching assignment of classes on each execution of *k*-means. A large *F*-score corresponded to a strong encoding of the feature-set of interest in the EEG data.

### Multiway Canonical Correlation Analysis (MCCA)

EEG data is notorious for its low signal-to-noise ratio (SNR), which represent one of the core challenges when analysing this kind of data. One approach to improve the SNR is multiway canonical correlation analysis (MCCA), a tool that identifies EEG components that are most correlated across subjects. Under the assumption that a stimulus would produce consistent cortical responses across subjects in the same age-group, MCCA identifies such consistent responses accepting that they may originate from distinct sources (i.e., distinct topographical patterns) for different subjects. MCCA is an extension of canonical correlation analysis^74^ to the case of multiple (> 2) subjects. Given N multichannel datasets Y_i_ with size T x J_i_, 1 ≤ i ≤ N (time x channels), MCCA finds a linear transform W_i_ (sizes J_i_ x J_0_, where J_0_ < min(J_i_)_1≤i≤N_), which, when applied to the corresponding data matrices, aligns them to common coordinates and reveals shared patterns^45^. These patterns can be derived by summing the transformed data matrices as follows: 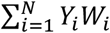. The columns of the matrix Y, which are mutually orthogonal, are referred to as summary components (SC). The first components are signals that most strongly reflect the shared information across the several input datasets, thus minimising subject-specific and channel-specific noise. Here, MCCA was run within each age group (4mo, 7mo, 11mo, and adults). After fitting the MCCA mapping and projecting the data to the SC space, a given number of component was retained (e.g., only the first component) before performing the inverse mapping and obtain a denoised version of the EEG signal for each subject. This denoising procedure was repeated by retaining a progressive number of components (**Figure 2B**).

### Statistical analysis

All statistical analyses directly comparing the groups were performed using a repeated measures ANOVA. One-sample Wilcoxon signed-rank tests were used for post hoc tests. Correction for multiple comparisons was applied where necessary via the false discovery rate (FDR) approach. In that case, the FDR adjusted *p*-value was reported. Descriptive statistics for the neurophysiology results are reported as a combination of mean and standard error (SE).

## Data and code availability

Analyses were conducted by using custom MATLAB scripts developed starting from publicly available scripts shared by the CNSP initiative (Cognition and Natural Sensory Processing; https://cnspworkshop.net). Such analysis scripts avail of external publicly available libraries: the mTRF-Toolbox (https://github.com/mickcrosse/mTRF-Toolbox)^8^, EEGLAB^68^; and the NoiseTools library (http://audition.ens.fr/adc/NoiseTools)^45^. Data was converted to the CND data structure (Continuous-event Neural Data - https://cnspworkshop.net), allowing to carry out the analyses with the CNSP analysis scripts, which provided a platform for bringing together all the necessary libraries. We commit to publicly share the EEG data in the first half of 2023. Study data were collected and managed using REDCap (Research Electronic Data Capture) electronic data capture tools hosted at Cambridge university^75,76^.

## Acknowledgements

We thank Dimitris Panayiotou, Alessia Philips, Natasha Mead, Helen Olawole-Scott, Panagiotis Boutris, Samuel Gibbon, Isabel Williams, Sheila Flanagan, and Christina Grey who helped collecting the data as well as all the families of the infant participants. We thank Dr. Susan Richards for her assistance on the phoneme transcription. We thank the CogHear workshop organisers (Mounya Elhilali, Malcolm Slaney, and Shihab Shamma) and participants for their useful feedback on the early results of this study.

## Footnotes

i This should not be intended as a hard boundary, as this is likely a gradual phenomenon that changes over large time windows, with differences between easy and more difficult speech contrasts

ii Please note that results did not change when acoustic-only TRFs consisted of acoustic vectors concatenated with shuffled phonetic information.

## Notes

Conflicts of interest: none declared.

Funding sources: This project received funding from the European Research Council (ERC) under the European Union’s Horizon 2020 research and innovation programme (Grant Agreement No. 694786) (A.A., U.G.). This research was conducted with the financial support of Science Foundation Ireland under Grant Agreement No. 13/RC/2106_P2 at the ADAPT SFI Research Centre at Trinity College Dublin (G.D.L., G.C.). ADAPT, the SFI Research Centre for AI-Driven Digital Content Technology, is funded by Science Foundation Ireland through the SFI Research Centres Programme. This work was also supported by the Science Foundation Ireland Career Development Award 15/CDA/3316 (G.D.L., R.R.). G.C. was supported by an Advanced European Research Council grant (NEUME, 787836) and by the FrontCog grant ANR-17-EURE-0017.

https://www.cne.psychol.cam.ac.uk/babyrhythm-project

https://cnspworkshop.net/resources.html

### Competing Interest Statement

The authors have declared no competing interest.

